# A matrix-mimicking bioadhesive epicardium for tunable modulation of biomechanics in the acutely infarcted heart

**DOI:** 10.1101/2025.10.07.681040

**Authors:** Claudia E. Varela, Diego A. Quevedo-Moreno, Jean Bonnemain, Keegan L. Mendez, Yiling Fan, Jonathan Tagoe, Carly Long, Mossab Saeed, Hyunwoo Yuk, Xuanhe Zhao, Ellen T. Roche

**Affiliations:** Institute for Medical Engineering and Science, Massachusetts Institute of Technology, Cambridge, MA, USA; Harvard-MIT Program in Health Sciences and Technology, Cambridge, MA, USA; Department of Mechatronics Engineering, Tecnológico de Monterrey, Monterrey, NL, MEX; Department of Mechanical Engineering, Massachusetts Institute of Technology, Cambridge, MA, USA; Department of Adult Intensive Care Medicine, Lausanne University Hospital and University of Lausanne, Lausanne, Switzerland; Department of Cardiac Surgery, Boston Children’s Hospital, Harvard Medical School, Boston, MA, USA; Department of Civil and Environmental Engineering, Massachusetts Institute of Technology, Cambridge, MA, USA

## Abstract

Mitigating adverse tissue remodeling after a heart attack or myocardial infarction (MI) is critical to prevent the development of heart failure. Among various post-MI treatment strategies, mechanical reinforcement of the infarcted region with epicardial patches has promise due to its consistent improvement of chronic cardiac function and its drug- or biologic-free nature. However, despite the variety of patch materials studied to date, the lack of a programmable platform that predictably modifies early-stage cardiac biomechanics to different degrees has prevented further optimization of this strategy. Here, we introduce the matrix-mimicking bioadhesive epicardium (MMBE), a platform that can be rationally designed to achieve a wide range of anisotropic mechanical properties to offer quantifiable mechanical reinforcement of the heart upon application. The platform synergistically combines fully programmable direct-ink-writing of extracellular matrix-inspired crimped fibers and a bioadhesive for sutureless integration to the epicardium. The MMBE platform achieves an array of matrix-mimicking mechanical properties and acute modulation of cardiac biomechanics using numerical analysis, in silico studies and experimental characterizations. Furthermore, the feasibility of the MMBE platform in an *in vivo* rat model of MI is demonstrated. The MMBE platform can be used to systematically identify patch design parameters that alter post-MI remodeling without introducing confounding biological variables.

## Introduction

After a myocardial infarction (MI), the infarcted muscle stops contracting and is stretched by adjacent, unaffected muscle negatively impacting the ventricle’s pumping capacity. Coupled to corresponding biological processes, these akinesis/dyskinesis contribute to the onset of adverse remodeling that can be conducive to heart failure^1,2^. Despite current clinical interventions, such as revascularization and medication, the risk of heart failure development in MI survivors remains considerable, which has prompted the development of additional interventions to mitigate this transition ^3,4^. Delivering cells, small molecules and other biological agents directly to the epicardial surface of the infarcted heart has proven to positively impact maladaptive remodeling after a MI in both preclinical and clinical settings^5–10.^

Another strategy that has consistently reduced characteristic late-stage adverse remodeling in preclinical models of MI consists of epicardially implanting biologic-free patches or devices with the goal of mechanically reinforcing or restraining the infarct region^11–13^. Implantation of epicardial patch materials such as hydrogels, decellularized scaffolds, polymers, and textiles (knitted or woven) has yielded a reduction in adverse remodeling and functional decay in small and large animal models ^10,12–19^. Although the biologic and drug free nature of this strategy circumvents many of the challenges associated with biologic interventions such as bioavailability, high production cost, toxicity, and inter-species response variability, there remains a lack of mechanistic understanding of how integrating inert patch materials to the infarcted heart results in chronic benefits.

Interestingly, chronic benefits are observed regardless of the implanted patch material and the effective reinforcement level the patch provides, as determined by the material mechanical properties and the level of tension imparted by sutures or adhesion-based attachment to the heart. Despite promising results achieved with a variety of materials and reinforcement levels, how specific patch design parameters contribute to acute and chronic improvements after MI remains elusive, complicating the advancement of this strategy and hindering its potential clinical impact. In fact, studies that characterize the acute biomechanical impact caused by these favorable epicardial patch materials are mostly lacking despite the consequential role that early-stage modification of infarct and ventricular biomechanics likely plays in subsequent remodeling.

To systematically investigate the biomechanical impact that reinforcing patches have in early post-MI adaptations and determine contributors to their late-stage benefits, a material platform that provides wide-ranging levels of reinforcement and predictably modulates cardiac biomechanics is needed. A suitable material platform should achieve a broad range of anisotropic mechanical tunability, informed by axis-specific behaviors from previously tested materials, without modifications to its chemical formulation to ensure comparable host biological responses regardless of the mechanical reinforcement provided. Also, the material platform must be easily customized and coupled to the heart of different animal models and produce rapidly quantifiable biomechanical changes upon implantation.

Here, we present the matrix-mimicking bioadhesive epicardium (MMBE), a programmable patch platform that can achieve wide-ranging anisotropic mechanical properties and produces biomechanical changes after adhering to the heart (Fig. 1a). The MMBE consists of an extracellular matrix-inspired crimped (i.e. sinusoidal) fiber mesh made by direct-ink-writing (DIW) of a hydrophilic polyurethane (PU) ink and a bioadhesive hydrogel^20,21^ for seamless coupling to the surface of the heart (Fig. 1a). The MMBE can be programmed to exert different levels of infarct reinforcement by customizing the morphology of the crimped fiber mesh which alters its strain hardening properties without changing its material composition (Fig. 1a). MMBEs developed to exert different levels of longitudinal restrain were found to proportionally decrease longitudinal strain in hydrogel tissue analogs under uniaxial and biaxial tension as well as in an *in silico* left ventricle (LV) model. To demonstrate the feasibility of the platform, the MMBE was implanted in a rat model of acute MI, leading to quantifiable cardiac biomechanical changes after infarction. MMBE designs that provide longitudinal reinforcement were the focus of this work to ensure a quantifiable acute cardiac response^17–19^. However, MMBE designs with different levels of anisotropic or isotropic infarct reinforcement can be produced for future studies.

**Figure 1.**
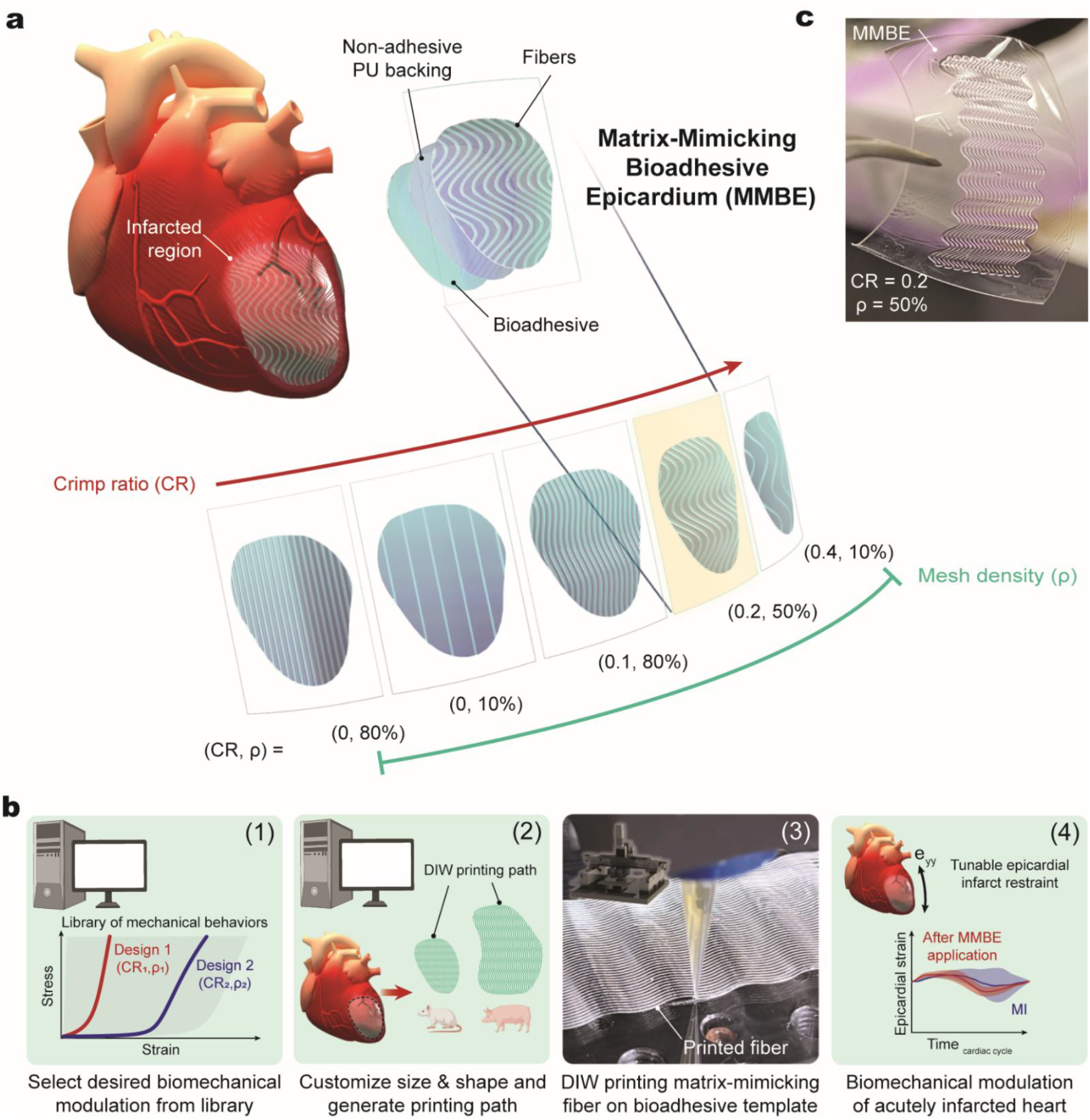
**a)** Schematic of the Matrix-Mimicking Bioadhesive Epicardium (MMBE) platform. The MMBE combines a bioadhesive hydrogel with direct ink writing (DIW)-printed fibers via a non-adhesive polyurethane (PU) backing for seamless integration to the infarcted heart epicardium. The extracellular matrix-inspired fiber mesh yields a wide range of MMBE designs by customizing parameters, such as crimp ratio (CR) and mesh density (ρ), to predictably modulate infarct biomechanics without altering the MMBE’s material chemistry. **b)** MMBE fabrication workflow. The desired MMBE uniaxial tensile behavior is selected from a model-generated library. After the size and shape of the MMBE are input, a custom-MATLAB program generates the required fiber mesh printing path (g-code). A DIW printer extrudes the desired fiber mesh design onto a prefabricated PU-coated bioadhesive template film. The MMBE can be directly implanted epicardially to acutely modulate infarct biomechanics. **c)** Image of a rectangular MMBE (CR=0.2, ρ=50%) fabricated with the developed workflow.

## Results

### Design and fabrication of MMBE platform

The MMBE platform is composed of a DIW-printed PU crimped fiber mesh integrated to a bioadhesive hydrogel^20,21^ via a non-adhesive PU backing (Fig. 1a). To allow for user-specific customization, the desired MMBE size, shape, and mechanical behavior/reinforcement profile are selected at the time of fabrication (Fig. 1b). First, the desired mechanical behavior is selected from a library of model-generated uniaxial tensile properties, yielding a unique combination of mesh design parameters *(crimp ratio, fiber thickness, and mesh density)*. Then, the desired MMBE size and shape are input into a custom-MATLAB program that generates the fiber mesh DIW printing path. Finally, a DIW-printer extrudes the fiber mesh with the desired characteristics onto a PU coated bioadhesive template film that can be directly implanted onto the epicardial surface of the heart (Fig. 1b-c).

### Mechanical tunability of MMBE platform enabled by DIW-printed mesh morphology

With this workflow, a wide range of mechanical properties can be achieved without altering the material composition of the MMBE platform, ensuring different MMBE designs elicit a comparable foreign body response after implantation. To this end, we incorporated an extracellular matrix (ECM)-inspired, crimped fiber mesh made of a biocompatible PU ink with high tensile strength whose morphology can be customized in terms of crimp ratio, fiber density, mesh and fiber thickness to program different uniaxial tensile behaviors in each MMBE mesh layer (Fig. 2a). Like many ECM proteins, such as collagen^22–24^, the crimped MMBE mesh is strain-stiffening, enabling us to toggle between different effective elastic moduli by delaying the onset of reinforcement. As the MMBE is stretched, a nonlinear J-shaped stress-strain curve is obtained (Fig. 2b). First, the curve is characterized by a bending-dominated behavior, where the thin crimped fibers unbend under relatively low magnitude forces. Once the fibers are fully straightened and elongation continues past this critical strain (ε_cr_), the behavior becomes stretch-dominated, where the force required to stretch the fully straightened crimped fibers increases relative to the elastic moduli of the PU ink, yielding a higher effective stiffness.

Finite Element (FE) simulations (Fig. 2b) were used to determine the uniaxial tensile behaviors that can be programmed to the MMBE platform based on the crimp ratio, fiber density, and fiber diameter. Increasing the crimp ratio allows us to control the onset of restraint without changing the number of fibers that comprise the MMBE mesh (Fig. 2c). By varying the crimp ratio of the ECM-inspired fibers the effective elastic moduli of the MMBE (1.3 kPa-2.8 MPa) can be reduced to about 2000 times of the hydrophilic PU ink’s moduli when the MMBE is stretched by 50% of its original length (Fig. 2c). Our model shows that increasing the density (Fig. 2d) of the MMBE fiber mesh yields designs of increasing overall stiffness, while varying the diameter of individual fibers results in MMBEs with relatively comparable mechanical properties (Fig. 2e).

To validate the simulated MMBE mechanical behaviors, MMBE meshes with matching features were fabricated and subjected to mechanical testing. Microscopy images of fabricated MMBE designs were used to verify the desired features were appropriately printed (Fig. 2f). Under uniaxial tensile loading (Fig. 2g), the measured behavior of MMBE fiber meshes with varying crimp ratios (Fig. 2h) and mesh densities (Fig. 2i) closely resembled the behavior predicted by our computational model.

### In silico reinforcement of infarcted LV model with MMBE platform

To evaluate how MMBEs with different reinforcement capabilities impacted LV muscle biomechanics, we developed a FE computational model of a porcine LV with a large anteroapical infarct integrated into a lumped parameter model of the circulatory system that reproduced functional parameters reported in literature^17,18,25^ (Fig. 3a, Supp. Fig. 1). MMBEs providing low, medium, and high degrees of longitudinal reinforcement were epicardially coupled onto the modeled LV ensuring coverage of the infarct and border zone (BZ) regions (Fig. 3b). To model the longitudinal and circumferential material properties of each MMBE design, the material coefficients required for the constitutive Holzapfel–Gasser–Ogden (HGO) hyperelastic anisotropic material model^26^ were calculated from uniaxial tensile data of the bioadhesive hydrogel for the circumferential axis and the J-shaped stress-strain data corresponding to the desired level of reinforcement in the longitudinal axis (Fig. 3c). FE analysis of selected MMBE designs yielded comparable axis-specific behaviors to experimental tensile data and their corresponding analytical HGO model fit, validating the HGO model is suitable for MMBE modeling (Fig. 3c).

**Figure 3.**
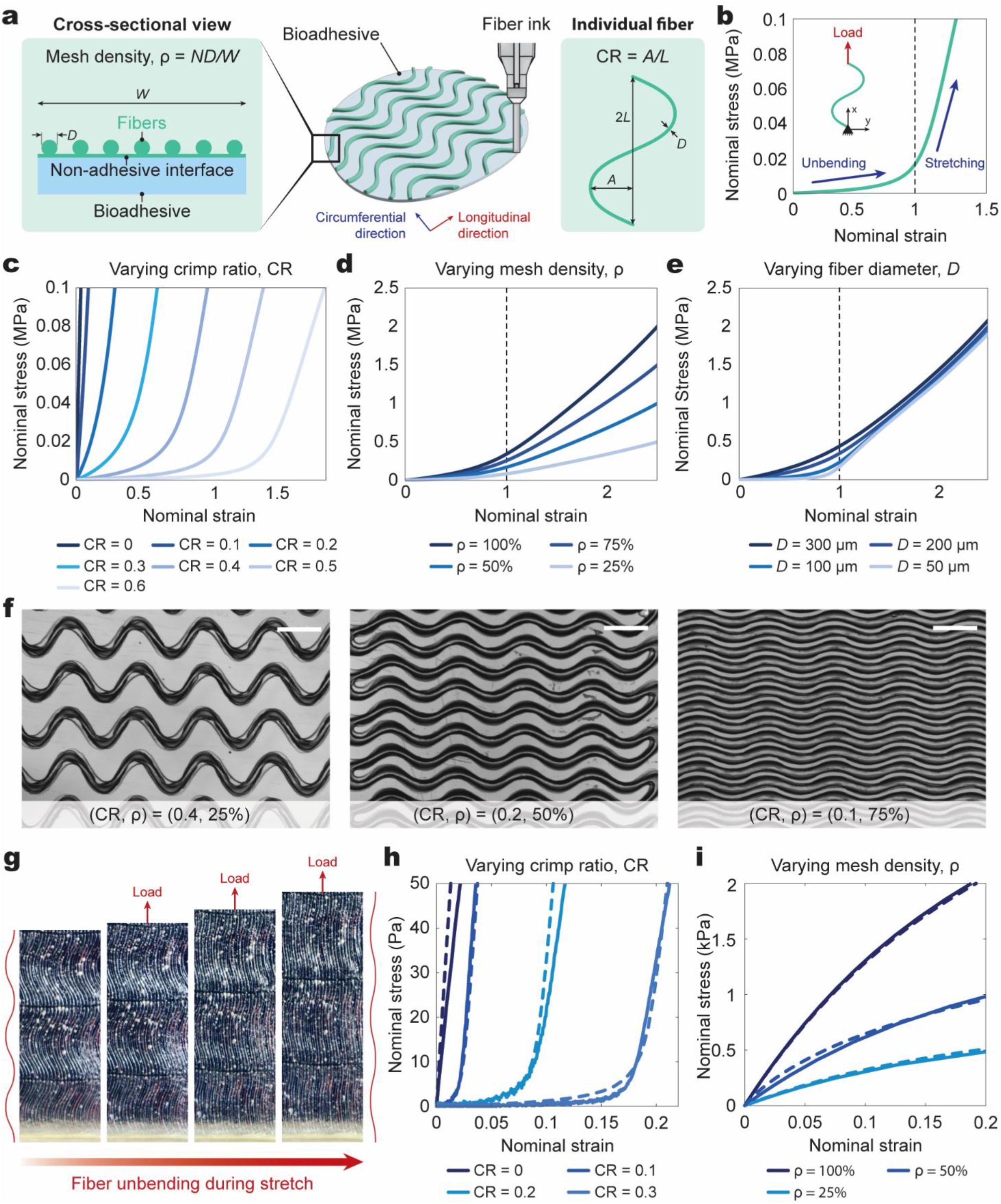
**a)** Cross-section of the MMBE platform components. A pre-determined number (N) of individual fibers of diameter (D) are extruded along the longitudinal direction to achieve the desired mesh density (ρ) across the MMBE’s width (W). Each fiber’s morphology is determined by its crimp ratio (CR), defined as the ratio of the fiber’s sinusoidal amplitude (A) and half a period (L). **b)** Like many extracellular matrix proteins, the MMBE sinusoidal fibers are strain-stiffening. Under a uniaxial tensile load, a J-shaped stress-strain curve is obtained. First, the sinusoidal fibers unbend until a critical strain (dashed line) is reached at which the fibers are fully straightened. Elongation beyond the critical strain becomes stretch-dominated; where the force required to continue stretching the now straightened fibers increases relative to the elastic moduli of the fiber ink. **c)** Simulated uniaxial tensile behaviors achieved by modifying the CR (0-0.6) of the MMBE fiber mesh. **d)** Simulated uniaxial tensile behaviors achieved by modifying the ρ (25-100%) of the MMBE fiber mesh. **e)** Simulated uniaxial tensile behaviors achieved by modifying D (50-300 um) of individual fibers in the MMBE mesh. **f)** Sample microscope images of various MMBE designs. **g)** Images of a MMBE fiber mesh as it unbends under uniaxial tensile loading. Scale bar=400 μm. **h)** Simulated (dashed line) and measured (solid line) behaviors of fabricated MMBE designs of varying CR. **i)** Simulated (dashed line) and measured (solid line) behaviors of fabricated MMBE designs of varying ρ.

Next, we assessed how selected MMBE designs impacted ventricular mechanics in our model. As previously reported^17,18^, an acute MI increases ventricular wall strain at the MI and BZ at the end of diastole (i.e. relaxation) (Fig. 3d) and introduces a high stress concentration zone at the edges of the infarct and BZ regions at the end of systole (i.e. contraction) (Fig. 3e). Upon coupling a high reinforcement MMBE, a reduction in fiber strain and stress in the infarcted region is observed, demonstrating that the MMBE is effectively reinforcing the zone (Fig. 3d-e). Attachment of low and medium reinforcement MMBEs yield different degrees of diastolic strain and systolic stress reduction when compared to the no MMBE condition (Fig. 3f-g). The average fiber strain (Fig. 3f) and stress (Fig. 3g) in the MI and BZ regions decrease as the level of reinforcement provided by the MMBEs increases.

### Anisotropic reinforcement of a tissue analog with MMBE platform

To validate that the MMBE platform can also modulate the mechanical behavior of a tissue analog, MMBEs of increasing longitudinal stiffness, achieved by manipulating the thickness of the MMBE fiber mesh, were coupled to rectangular polyacrylic acid (PAA)-gelatin hydrogels and subjected to uniaxial tensile testing (Supp. Fig. 2a). The longitudinal tensile strength of the tissue analog increased as stiffer MMBE designs were coupled when compared to the tissue analog without an MMBE attached (Supp. Fig. 2b), confirming that modifying the longitudinal stiffness of the MMBEs fiber mesh directly modifies the longitudinal behavior of the tissue analog.

Then, we characterized how attaching MMBEs with different stiffness and strain-hardening properties modified the biaxial mechanical behavior of a tissue analog. MMBEs were coupled to PAA-gelatin hydrogels for biaxial loading using a pressurized chamber set up (Fig.4a-b). The circumferential (E_xx_) and longitudinal (E_yy_) strain experienced by the tissue proxies upon one pressurization cycle were calculated using digital image correlation. Coupling an MMBE with higher longitudinal stiffness (design 3, *ρ*=75%) to the tissue analog yielded a moderately greater reduction in longitudinal strain than the reduction achieved by an MMBE of lower stiffness (design 1, *ρ*=25%) with respect to the case where no MMBE was attached (Fig. 4c-g).

**Figure 4.**
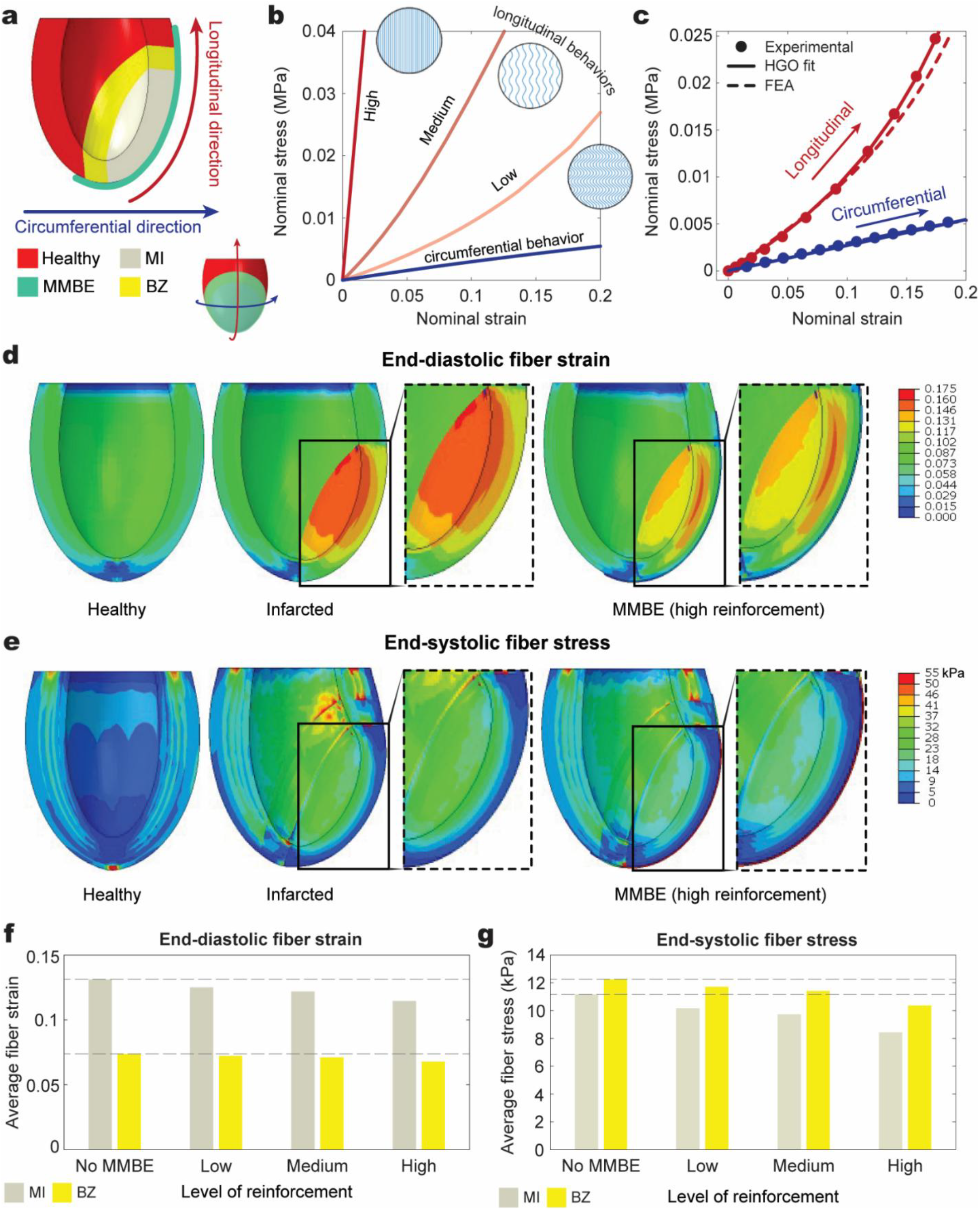
**a)** Cross-section of the simulated left ventricle (LV) with a large anteroapical myocardial infarct (MI) and border zone (BZ) covered by a Matrix-Mimicking Bioadhesive Epicardium (MMBE). **b)** Simulated longitudinal tensile behaviors of the tested MMBE designs providing low, medium, and high degrees of longitudinal reinforcement. The circumferential tensile behavior of all MMBEs is determined by the measured properties of the bioadhesive hydrogel film. **c)** Representative curves showing correspondence between the desired longitudinal and circumferential MMBE behaviors (dots), the calculated Holzapfel–Gasser–Ogden (HGO) constitutive model behaviors (solid line), and the MMBE behaviors obtained (dashed line) after LV filling via finite element analysis (FEA). **d)** End-diastolic strain of a healthy, infarcted, and MMBE-reinforced LV. **e)** End-systolic fiber stress of a healthy, infarcted, and MMBE-reinforced LV. **f)** Bar graph of averaged end-diastolic fiber strain across the MI and BZ regions in an LV with no MMBE (infarcted), a low, medium, and high reinforcement MMBE. **g)** Bar graph of averaged end-systolic fiber stress across the MI and BZ regions in an LV with no MMBE (infarcted), a low, medium, and high reinforcement MMBE.

Apart from modulating the magnitude of longitudinal reinforcement by changing the MMBEs mesh density, we controlled the onset of longitudinal restraint by increasing the amplitude of the ECM-inspired fiber mesh (design 2, CR=0.5). The immediate onset of longitudinal restraint achieved by the MMBEs with a crimp ratio of zero (design 1 and design 3) is observed by the marked reduction in longitudinal strain in the region of the tissue analog covered by the MMBE (Fig. 4c-g). An MMBE with a crimp ratio larger than zero (design 2) achieved a delayed onset of restraint as evidenced by the minimal changes in longitudinal strain when compared to the tissue analog without an MMBE attached (Fig. 4c-g).

Notably, the magnitude of circumferential strain (E_xx_) remains largely unaffected regardless of the coupled MMBE design, confirming the MMBEs attached exclusively provide the intended anisotropic longitudinal restraint (Supp. Fig. 3a). Longitudinal and circumferential strain were unaffected when only the bioadhesive layer of the MMBE platform (i.e. no fiber mesh) was coupled to the tissue analog (Supp. Fig. 3b-e), confirming that the DIW-printed, ECM-inspired fiber mesh is responsible for the anisotropic restraint capabilities of the MMBE platform.

### In vivo modulation of infarct biomechanics with the MMBE platform

After validating that our MMBE platform imparts a variety of reinforcement regimes without altering its material composition, the ability of the MMBE platform to elicit quantifiable biomechanical changes in a terminal rodent MI model of left anterior descending (LAD) coronary artery occlusion was investigated. MMBE designs validated to provide immediate longitudinal restraint *in vitro* (Fig. 4b: design 1 (CR=0, *ρ*=25%) and design 3 (CR=0, *ρ*=75%)) were attached to the infarct region 30 min after LAD ligation.

Healthy heart muscle shortens in a coordinated manner during systole (yielding increasingly negative longitudinal strain values) and relaxes back to its original length during diastole (seen as decreasing longitudinal strain until a value of zero is reached) (Fig. 5a). Conversely, acutely infarcted muscle is unable to contract and instead becomes stretched during systole and recoils during diastole (Fig. 5a). To assess the biomechanical impact of selected MMBEs, the longitudinal epicardial strain of 10 points along the infarcted LV anterior wall was quantified before and after MMBE placement using speckle tracking of long axis view (LAX) echocardiography videos of at least one cardiac cycle (Fig. 5b). Two experienced echocardiogram readers identified if the individual points were in a muscle region covered by the attached MMBE and if the contractility of the muscle region was normal (healthy, H), reduced (border zone, BZ), or non-existent (infarcted, MI) (Fig. 5b-e; Supp. Fig. 4, Supp. Fig. 5). Only animal subjects in which MMBEs covered >50% of MI area were included in the study groups.

**Figure 5.**
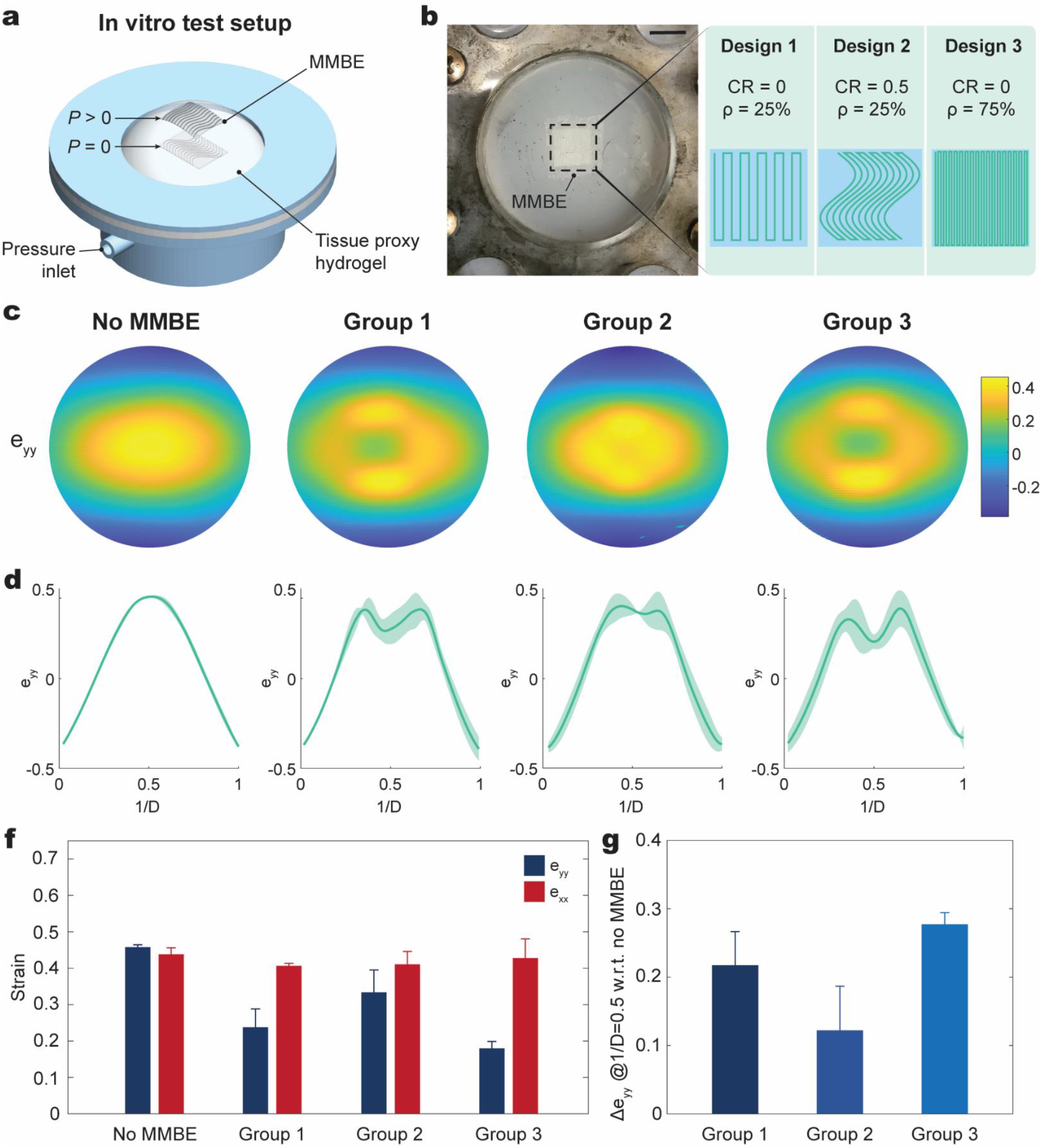
**a)** Schematic of pressurized chamber set up used for MMBE biaxial loading. Pressurization (P>0) of the chamber stretched the MMBE-reinforced polyacrylic acid hydrogel (tissue proxy). **b)** Image of an MMBE coupled to the tissue proxy along the MMBE designs used in each experimental group. Scale bar= 7 mm. **c)** Representative longitudinal (eyy) strain heat map at 18 mmHg of chamber pressure calculated from digital image correlation (DIC). **d**) Longitudinal strain (eyy) values across the horizontal centerline of the recorded picture frames of width D. **f)** Bar graph of averaged strain values (eyy; exx) per experimental group. **g)** Bar graph of calculated difference of longitudinal strain (eyy) at the center point of the picture frame in the no MMBE group and each experimental group. n=3 per group; data reported as mean± st.dev.

The longitudinal strain curves obtained from each point on the muscle region attached to the MMBEs (i.e. treatment region) (Fig. 5c, e) were used to calculate the mean longitudinal strain and standard deviation during one cardiac cycle (Fig. 5c, e; Supp. Fig. 4, Supp. Fig. 5). The longitudinal strain variability, measured by calculating the area spanning one standard deviation about the mean longitudinal strain, significantly decreased after placement of MMBE design 1 (Fig. 5g, Supp. Fig. 4) and followed a similar trend after placement of MMBE design 3 (Fig. 5i, Supp. Fig. 5). This quantifiable reduction of longitudinal muscle motion on the infarcted wall indicates that attachment of the selected MMBEs imparts a measurable degree of longitudinal restraint on the acutely infarcted heart.

## Outlook/Discussion

This work introduces the MMBE, a rationally designed patch platform capable of diverse and quantifiable biomechanical modulation of the infarcted heart. The MMBE is composed of a biocompatible DIW-printed fiber mesh and a bioadhesive hydrogel for sutureless integration to the epicardial surface of the heart. The MMBE fiber mesh can mimic the strain hardening behavior of ECM proteins, like collagen, and allows the user to program a wide range of anisotropic mechanical properties without altering the material building blocks of the MMBE platform (Supp. Fig. 6a). Numerical analysis and mechanical testing were used to map the scope of mechanical tunability that can be programmed onto the MMBE. The biomechanical modulation capability of the MMBE was demonstrated by the tunable longitudinal restraint provided to a porcine LV *in silico* and to tissue proxies under uniaxial and biaxial load. Finally, the feasibility of MMBE biomechanical modulation *in vivo* was demonstrated in a terminal rat model of acute MI. Coupling MMBEs designed to restrain aberrant motion in the longitudinal axis of the heart led to a reduction in strain variability of the patched region.

**Figure 6.**
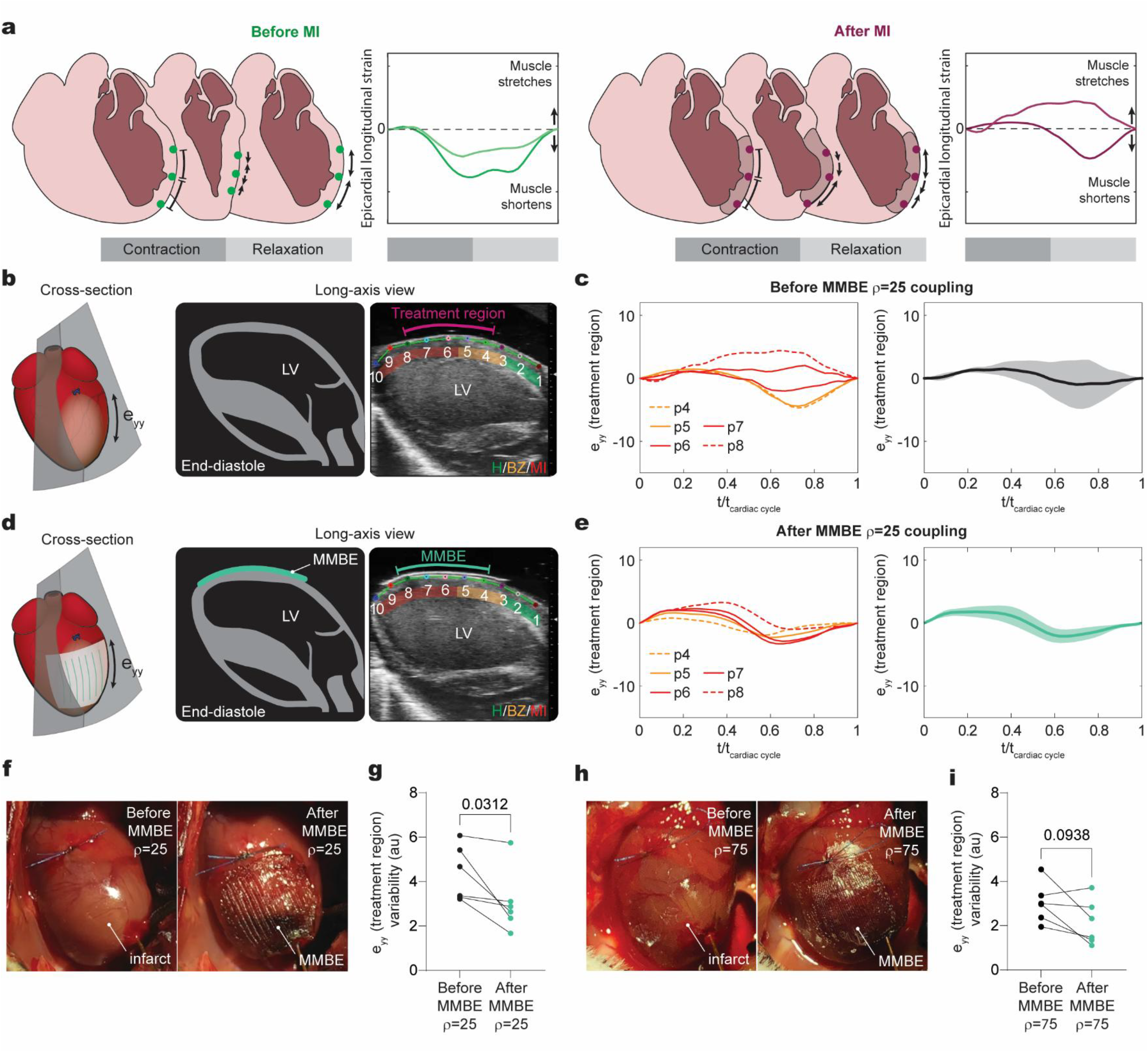
**a)** Schematic of longitudinal muscle motion in infarcted and non-infarcted muscle during one cardiac cycle. Schematic and sample image of long axis view (LAX) in the infarcted **(b)** and MMBE-reinforced (design 1; ρ=25) **(d)** heart obtained by echocardiography. 10 points were placed along the epicardium for longitudinal strain quantification in one cardiac cycle and classified to have normal (H), reduced (BZ), or non-existent (MI) muscle contraction and be within or out of the treatment region covered by the MMBE. The mean longitudinal (eyy) strain curve (± st.dev.) was calculated from individual strain curves per point inside the treatment region before **(c)** and after **(e)** MMBE reinforcement (design 1; ρ=25). **f)** Image of infarcted rat heart before and after MMBE reinforcement (design 1; ρ=25). **g)** Graph showing the longitudinal strain variability measured as the area spanning one standard deviation about the mean eyy curve of all subjects before and after MMBE placement (design 1; ρ=25) (n=6). **h)** Image of infarcted rat heart before and after MMBE reinforcement (design 3; ρ=75). **i)** Graph showing the longitudinal strain variability measured as the area spanning one standard deviation about the mean eyy curve of all subjects before and after MMBE placement (design 3; ρ=75) (n=6).

The MMBE platform was developed to enable systematic studies that help identify the material design parameters that alter post-MI remodeling without introducing confounding variables associated with material customization. Chronic cardiac improvements after implantation of acellular materials have been reported, but a mechanistic link between specific features of the implanted materials and cardiac remodeling cannot be extrapolated because a wide range of material compositions, mechanical properties, and reinforcing capabilities seem to yield comparable chronic effects^10,12–19^. A few studies have explored how altering the mechanical properties of implanted epicardial materials through material composition changes results in different degrees of chronic functional recovery ^15,27^. The MMBE platform enables a more encompassing range of mechanical behaviors simply by modifying the MMBE’s fiber mesh morphology.

The components and fabrication workflow used to realize the MMBE not only allow for ample customization for implantation in animal models but can also be compatible with clinical settings. The mechanical behavior, size and shape of the MMBE can be personalized at the time of fabrication based on the ink utilized, the mesh morphology, and the characteristics of the infarct region onto a pre-fabricated bioadhesive template that can be cut to size. The range of tensile strength achieved by the MMBE platform was guided by the elastic moduli of previously reported patch materials although a DIW-ink of higher tensile strength would be required for an MMBE design that mimics the highest elastic moduli reported^17,19^. While the current work focused on validating the biomechanical modulation capabilities of PU-based, longitudinally reinforcing MMBEs of rectangular shape and 7 mm^2^ size, larger MMBEs with other levels of reinforcement can be generated (Supp. Fig. 6b), making the MMBE platform suitable for both small and large animal models. Some platform features that may facilitate the clinical adoption of the MMBE include the ability to produce MMBEs of infarct-specific shapes segmented from clinical imaging modalities such as MRI (Supp. Fig. 6c) as well as the adhesive and atraumatic nature of MMBE delivery that can be compatible with less invasive implantation procedures.

To assess whether the observed biomechanical effects reflect the designed mechanical properties of the MMBEs, we quantified the acute *in vivo* response following the application of two longitudinally restraining MMBEs to the heart. The implanted MMBE designs were selected for these feasibility studies because acute longitudinal restraint of infarcted dog hearts has been shown to produce a quantifiable biomechanical response 30 min after reinforcement^17,18^. The MMBEs selected for implantation were first shown to reduce the magnitude of longitudinal strain of a tissue analog to moderately different degrees (Fig. 4f-g) prior to implantation in a rat model. Significant longitudinal restraint of the rat heart was achieved with MMBEs design 1 (Fig. 5g). Interestingly, the heart seems to respond similarly when a higher effective stiffness MMBE design (design 3) is coupled (Fig. 5i), which may indicate that the degree of stiffening provided by design 1 is sufficient to produce a quantifiable response or the level of increased stiffness of design 3 is not enough to elicit a different biomechanical response within the studied timeframe.

From this initial *in vivo* feasibility assessment, we learned that decreasing variability in MMBE placement with respect to the infarct size and shape is an important consideration for future chronic studies. Despite of consistently located LAD ligations and MMBE placement on top of the blanched ventricular area observed following LAD ligation, the amount of infarcted and hypocontractile BZ muscle covered by the MMBE was different among animals as indicated by the echocardiogram scoring (Supp. Fig.5b, 6b). Although these variations limited the analysis to paired changes in longitudinal strain variance within the treatment region of each animal, the data indicates that implantation of the MMBE modulated the biomechanics of the patched region, thus validating the potential utility of the MMBE platform. To circumvent the observed variation in future rodent studies a larger MMBE can be implanted to ensure complete coverage of the infarcted and hypocontractile regions. Proper MMBE placement could be confirmed by echocardiogram at the time of implantation and corrected if needed since the bioadhesive layer can be detached on demand^28^.

Although the MMBE components are biocompatible, have long lasting efficacy and favorable foreign body responses^21^, future chronic studies will investigate the MMBE’s platform potential for sustained biomechanical modulation, integration and mitigation of adverse remodeling and functional decay. It is possible that the surface level coupling enabled by the MMBE’s bioadhesive may overcome the unloading observed in other acellular patches attached to the heart by sutures as the infarct scar compacts weeks post-MI^19^. Finally, without introducing confounding biological variables associated with alterations in material chemistry, the MMBE platform will enable systematic investigations to help identify specific patch design parameters and mechanical reinforcement regimes that alter post-MI remodeling, holding significant promise for accelerating the advancement of this biologic and drug-free strategy and enhancing its potential clinical impact.

## Materials and Methods

### MMBE Fabrication

#### Bioadhesive template

As previously reported^20,21^, to prepare the bioadhesive hydrogel, a stock solution containing acrylic acid (30 w/w%), chitosan (Sigma 95/500, 2 w/w%), *α*-ketoglutaric acid (0.2 w/w %), and poly(ethylene glycol methacrylate) (PEGDMA; Mn = 550, 0.03 w/w%) were dissolved in deionized water. Then, 50 mg of N-acryolsuccimide (Acros Organics B.V.B.A.) were dissolved to 10 ml of the above stock solution to get a precursor solution. The precursor solution poured on a glass mold with 210 *μ*m-thick spacers and cured in an ultraviolet light chamber (354-365 nm, 10-12 W power) for 30 min. The non-adhesive layer, 10 w/w% hydrophilic polyurethane (PU) (AdvanSource Biomaterials) in ethanol/water mixture (95:5 v/v) was spin-coated on the bioadhesive at 400 rpm for 60 s. Inside a biosafety hood, the bioadhesive template was stretched in both the length and width direction based on a previously reported method^29^ to limit swelling to the thickness direction and dried under air flow for at least 1 hour.

#### MMBE fiber mesh

The crimped (i.e. sinusoidal) MMBE fiber mesh was printed using a fiber ink made of hydrophilic PU (HydroThane, AdvanSource Biomaterials) in 50% dimethylforamide (DMF) & 50% tetrahydrofuran (THF) at a concentration of 30 w/v%. A unique combination of mesh design parameters *(crimp ratio, fiber thickness, and mesh density)* and the desired MMBE size and shape are input into a custom-MATLAB program that generates the desired fiber mesh DIW printing path (i.e. g-code). The desired MMBE mesh was directly printed onto a dry PU-coated bioadhesive layer secured onto a glass substrate. A custom-built direct-ink writing printer was used to print all MMBEs (Soft Active Materials Lab, MIT). Printing speed, extrusion pressure, and nozzle diameter were adjusted to extrude fibers of 150-200 *μ*m in diameter for all rodent MMBEs and 400-500 *μ*m in diameter for the MMBE adhered onto a porcine heart (CR=0.5, r=25%). To achieve the desired fiber mesh thickness (0.1 mm for rodent MMBEs and 0.45 mm for the porcine MMBE), the required number of layers to be extruded was calculated from single layer measured thickness. Fully assembled MMBEs were sealed in a plastic bag with desiccants (silica gel packets with active charcoal, McMaster Carr) and stored at –20 ℃ before use. Digital images of selected MMBEs were taken using a Nikon Eclipse LV100ND microscope.

To obtain a patient-specific MMBE size and shape, a deidentified cardiac MRI dataset from a late stage post-MI patient was processed to obtain a 3D mesh of the LV muscle using Materialise Mimics. The 3D meshes of the inner and outer LV wall were simplified and separated using SolidWorks. The thickness of the LV muscle was calculated using MATLAB by determining the distance between each point of the inner and outer wall meshes. Given the expected wall thinning in the area, the infarct region was defined by mesh distances less than 5mm, and the corresponding mesh elements were exported into MATLAB. To define the infarct shape for MMBE customization, a 2D projection of the infarct region’s outline was calculated using MATLAB and 5mm added to the perimeter to ensure that if printed, the patient-specific MMBE would be anchored to a non-infarcted area of the heart. Finally, an arbitrary combination of fiber mesh design parameters was used to produce a patient-specific shaped MMBE.

### Numerical simulations of MMBE’s mechanical behavior library

Numerical simulations were performed to determine the possible range of mechanical behaviors the MMBE platform can have under tensile load by varying MMBE design parameters: sinusoidal crimp ratio, fiber diameter, and relative mesh density. All numerical simulations were performed using commercial FE package ABAQUS (Dassault Systèmes, Providence, RI, USA). The MMBE platform was modeled as an anisotropic fiber-reinforced composite given that each MMBE design is composed of crimped (i.e. sinusoidal) fibers in series and in parallel coupled to a bioadhesive^30,31^. An individual or an array of longitudinally aligned crimped fibers were defined (Eq.1) and programmed to have the measured elastic moduli of the hydrophilic PU ink and bioadhesive in the longitudinal direction and circumferential direction, respectively. All models were discretized by meshing the geometries with linear hexahedral elements (C3D8) and accuracy was confirmed with a mesh refinement study. Geometry nonlinearity was implemented to simulate large structural deformations. Crimp ratio (CR), a single dimensionless parameter relating sinusoidal amplitude (A) and wavelength (L) was defined by Eq. 2. Relative fiber mesh density of the MMBE (*ρ*), where ***A***_***fiber***_ is the area of fiber and ***A***_***sample***_ is the area of sample was defined by Eq. 4 and determines the level of tensile strength of the MMBE (Eq. 5) where ***s*** is the MMBE strength and ***S*** is the fiber strength.

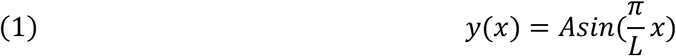

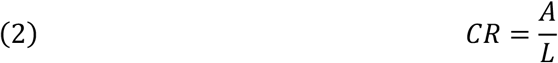

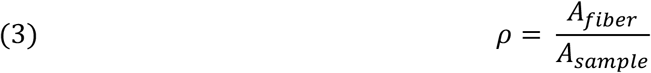

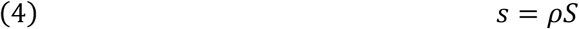

The critical strain of the stress-strain curve transition from bending-dominated to stretch-dominated behavior was calculated with Eq. 5.

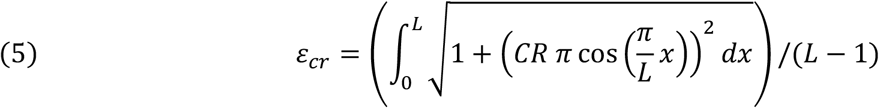

Numerical simulations of individual MMBE mesh fibers of varying CR were done with a 100 μm fiber diameter (*D*) and varying *D* simulations were done with a CR=0.5. Varying ρ simulations were done with a CR=0.5 and *D*=100 μm.

### Mechanical testing of MMBE fiber mesh designs

Uniaxial tensile testing of select MMBE fiber meshes was performed with a mechanical tester (20 N loadcell, Zwick/Roell Z2.5). All tests were conducted with a constant tensile speed of 50 mm/min. The measured load-displacement curves were used to calculate the nominal stress-strain curve based on the dimensions of the printed samples (width=5mm, length = 50mm and thickness =0.125mm) using MATLAB. Representative measured MMBE stress-strain curves were compared to the computationally predicted stress-strain curve of the corresponding MMBE design.

### In silico left ventricular porcine model

All simulations were conducted in Abaqus (Dassault Systèmes, Providence, RI, USA). A truncated ellipsoid was used to represent an idealized left ventricle of the heart^32^, and its dimensions were scaled to match previously reported end-diastolic volume for a healthy pig heart^25^. The LV geometry was partitioned layer by layer to set the cardiomyocyte arrangement from a fiber angle of 60⁰ at the endocardium to an angle of −60⁰ at the epicardium in accordance with the Holzapfel and Ogden model^33,34^. The LV was meshed with hexahedral elements (C3D8) and a mesh refinement study was conducted to ensure accuracy of the results.

The passive material behavior for cardiac tissue was modeled using a transversely isotropic hyperelastic model^35^ and implemented using the Fung strain energy density function (Eq. 6).

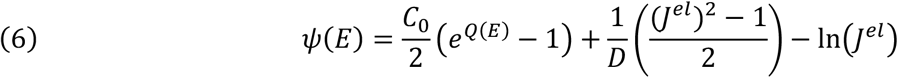

Since the myocardium has low compressibility, a high bulk modulus was defined, causing parameter *D* to be close to zero^36^. *J*^*el*^ is the elastic volume ratio. *Q*(*E*) is an exponent term function of the previously defined^35^ Green-Lagrange strain tensor ^35^in Eq. 7, where the subscripts *c,s* and *n* are the cardiomyocyte, sheetlet, and sheetlet-normal directions, respectively. The coefficients for the Eq. 7 were replicated from McGarvey et al^37^ and are shown in Table 1.

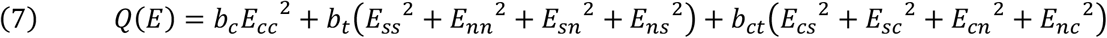

**Table 1.** Passive material parameters.

| Parameter | Definition | Value |
| --- | --- | --- |
| $C_0$ (kPa) | Elastic Constant | 0.39 |
| $b_c$ | Stiffness in fiber direction | 72.03 |
| $b_t$ | Stiffness in cross-fiber direction | 5.08 |
| $b_{ct}$ | Stiffness in shear direction | 36.69 |

The active strain approach was used to represent contraction of the cardiomyocyte fibers along its direction^38^ and commonly only depends on the fractional shortening of the heart^36^. A thermal strain tensor was introduced to simulate active myocyte contraction^36^, and is defined by Eq. 8, where *ε*_*th*_ is the thermal strain tensor, *α* is the thermal expansion coefficients and Δ*T* is the temperature difference.

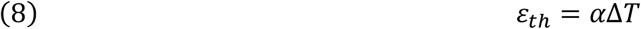

The thermal expansion coefficients were tuned to match previously reported healthy pressure-volume relationships^25^. The coefficient values used are shown in Table 2.

**Table 2.** Thermal expansion coefficients.

| Coefficient | Definition | Value |
| --- | --- | --- |
| $\alpha_{cc}$ | Cardiomyocyte direction | -0.16 |
| $\alpha_{ss}$ | Sheetlet direction | 0.105 |
| $\alpha_{nn}$ | sheetlet-normal direction | 0.105 |

The remaining tensor coefficients were set to zero to achieve an orthotropic strain where cardiac tissue contracts in the cardiomyocyte direction and expands in the sheetlet and sheetlet-normal directions^36^. To model a dynamic contraction in the ventricle, a curve based on the active tension of the cardiomyocytes ^39^ was input as a predefined temperature field to vary the temperature of the model and mimic the contraction according to the time of the cardiac cycle.

To model the hemodynamics of a full cardiac cycle, the LV was coupled into a lumped parameter model^40^ (Supp. Fig. 1a). This model was defined through a combination of surface-based fluid cavities and predefined fluid exchange properties^41^. The mass flow rate between two cavities is defined by Eq. 9. Where 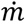 is the mass flow rate, *ρ* is the blood density, 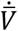 is volumetric flow rate per unit area and *A* is the effective exchange area. The mass flow of the exchange is related to the pressure difference (Δ*p*) between the cavities and to a viscous resistant coefficient (*C*_*v*_) (Eq. 10).

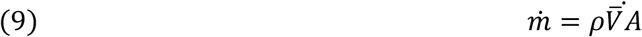

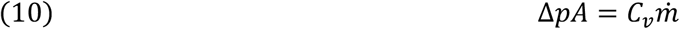

The lumped parameters used to represent the circulatory model were replicated from Sack et al^25^ and are shown in Table 3.

**Table 3.** Lumped Parameters of the circulatory system.

| Parameter | Definition | Value |
| --- | --- | --- |
| $R_A$ | Aortic valve resistance | 0.03 mmHg s ml<sup>-1</sup> |
| $R_M$ | mitral valve resistance | 0.03 mmHg s ml<sup>-1</sup> |
| $R_{Sys}$ | systemic arterial resistance | 1.7 mmHg s ml<sup>-1</sup> |
| $R_T$ | tricuspid resistance | 0.03 mmHg s ml<sup>-1</sup> |
| $R_P$ | pulmonary valve resistance | 0.03 mmHg s ml<sup>-1</sup> |
| $C_A$ | Systemic arterial compliance | 0.8ml mmHg<sup>-1</sup> |
| $C_V$ | venous compliance | 16.6ml mmHg<sup>-1</sup> |
| $C_P$ | Pulmonary compliance | 4.43ml mmHg<sup>-1</sup> |

A large anteroapical infarct was simulated by turning off the active contraction and changing the passive material properties in the infract region. To simulate the apparent softening^18^ in the acute infarct region, the stiffness of the passive material stiffness as determined by *C*_0_ was calibrated to match the percentage of volume increased after the infarction reported in literature^17^. A border zone (BZ)^42^ connecting the infarcted and healthy tissue was defined by conserving the passive material properties and decreasing the contractility by 50%^36^. A summary of the material properties in each zone is presented in Table 4.

**Table 4.**
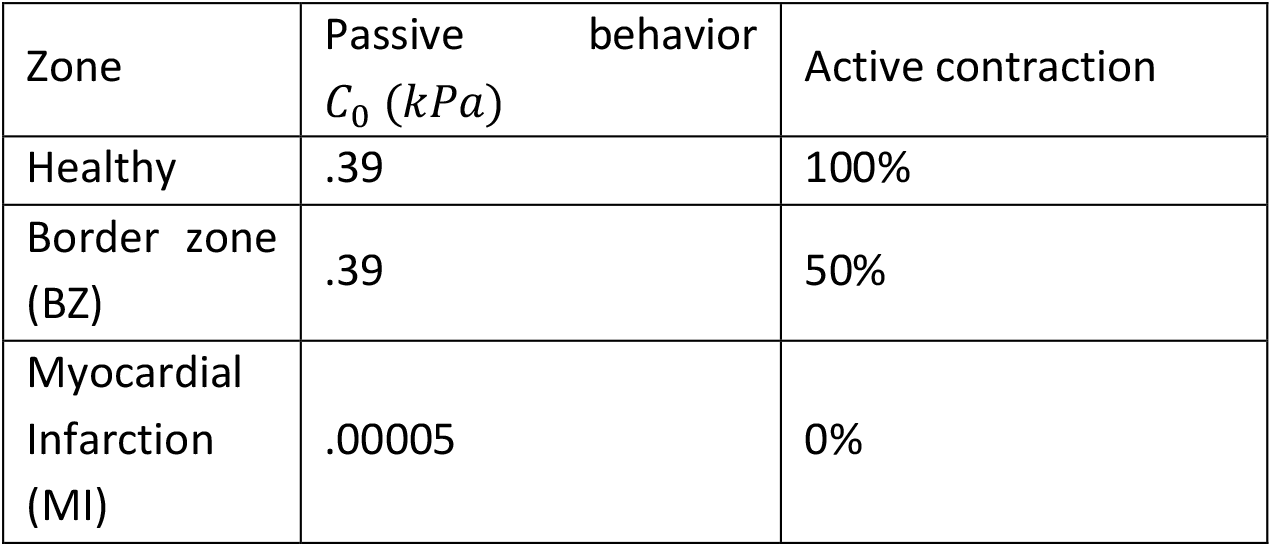
Change in the material properties to simulate a MI.

| Zone | Passive behavior<br>$C_0$ (kPa) | Active contraction |
| --- | --- | --- |
| Healthy | .39 | 100% |
| Border zone (BZ) | .39 | 50% |
| Myocardial Infarction (MI) | .00005 | 0% |

**Table 5.** MMBE designs and HGO material parameters.

| Reinforcement | MMBE Design Parameters |  |  | HGO Material Parameters |  |  |
| --- | --- | --- | --- | --- | --- | --- |
| | CR | $\rho$ (%) | t(mm) | $C_{10}$ | $k_1$ | $k_2$ |
| **Low** | 0.4 | 50 | 0.1 | 0.005 | 0.0204 | 0.2499 |
| **Medium** | 0.2 | 25 | 0.1 | 0.005 | 0.0552 | 0.00150 |
| **High** | 0 | 75 | 0.1 | 0.005 | 0.4372 | 1.47E-08 |

MMBEs coupled *in silico* were modeled using the constitutive Holzapfel–Gasser–Ogden (HGO) hyperelastic anisotropic material model^26^ which allows for simulation of the programmed J shaped stress-strain behavior in the longitudinal direction, while maintaining the mechanical properties of the adhesive hydrogel in the circumferential direction.

The Holzapfel–Gasser–Ogden strain energy function is defined by Eq. 11-12.

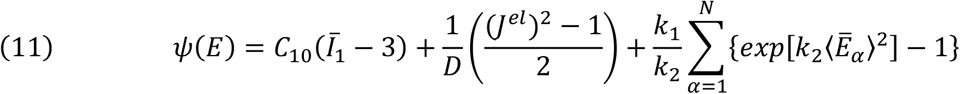

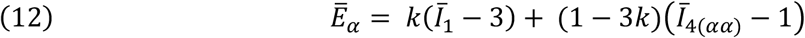

The model is defined by the constant material parameters ( *C*_10_, *D*, *k*_1_, *k*_2_ *and k*), where *C*_10_ and *D* define the matrix stiffness and matrix compressibility respectively, since the matrix of the MMBEs is assumed to be an incompressible material, *D* is set to be close to 0. *k*_1_ and *k*_2_ define the fiber non-linear behavior. And *k* defines the dispersion of the fibers in the matrix, in our case, the fibers are perfectly aligned to the longitudinal direction, so *k* is fixed to be 0. Then only 3 material constants need to be calculated to simulate any given MMBE design.

To determine the HGO coefficients (*C*_10_, *k*_1_, *k*_2_), needed to simulate a specific longitudinal or circumferential mechanical behavior, nonlinear least squares fit of the desired longitudinal and circumferential stress-strain curves were performed using MATLAB. The adhesive hydrogel uniaxial tensile data was used to determine the circumferential MMBE mechanical behavior in all simulations.

To study the effect that different levels of longitudinal reinforcement have in cardiac function, 3 different MMBE designs were selected from the model-derived library such that they provide a low, medium, and high level of longitudinal reinforcement. The corresponding design and HGO material parameters incorporated in the respective simulations are shown in Table 6 and a sample fit is shown in Fig. 3c. The modeled MMBEs were coupled into the LV covering the MI and BZ in all simulations.

### Mechanical testing of MMBE-reinforced tissue analogs

To make the PAA-gelatin hydrogel used as tissue analog, a stock solution containing 10 w/w% gelatin (300 Bloom Type A, Sigma), 20 w/w% acrylamide (Sigma) in DI water was made. Then, we dissolved 400 *μ*l 0.2M Ammonium persulfate (APS) solution, 400 *μ*l 0.23 w/w% N’,N’-Methylenebis(acrylamide) (MBAA) solution, and 10 *μ*l N,N,N’,N’-Tetramethylethylenediamine (TEMED) per 10mL of the stock solution to form the precursor solution. The precursor solution was then poured on a glass mold with 1.5 mm-spacers and cured for 1 h at room temperature.

MMBE designs of varying fiber mesh thickness (CR= 0.3; ρ=25%) were attached to rectangular tissue analogs and loaded onto a mechanical tester (20 N loadcell, Zwick/Roell Z2.5). Uniaxial testing of MMBE-reinforced tissue analogs was conducted at a constant tensile speed of 50 mm/min and the nominal stress-strain curves calculated using MATLAB.

For biaxial loading of MMBE-reinforced tissue analogs, a custom device was fabricated via 3D printing (Object). The tissue analog was screwed to the cylindrical chamber in between 2 acrylic plates laser cut to shape. The MMBE being tested was attached to the mounted tissue analog and marked with carbon powder for digital image correlation measurements. A light was placed under the tissue analog, on the bottom slot of the device, to generate sufficient image contrast. A syringe pump (Harvard Apparatus) was used to pressurize the chamber at a rate of 10 ml/min until a change of 18 mmHg was achieved. Recorded video frames from one pressurization cycle were imported into Vic 2D 2009 to calculate the circumferential (E_xx_) and longitudinal (E_yy_) strain of the tissue analog.

### in vivo strain measurements

Animal procedures were reviewed and approved according to ethical regulations by the Institutional Animal Care and Use Committee at Massachusetts Institute of Technology. Female Sprague Dawley rats (225-275 g) were used for these studies. Rats were anaesthetized using isoflurane (1-3% isoflurane in oxygen) in an anesthetizing chamber and their chest hair was removed. Endotracheal intubation was performed, and the rats were connected to a mechanical ventilator (Model 683, Harvard Apparatus) and placed over a heating pad for the duration of the surgery. A sternotomy was performed, and the pericardium was removed using forceps. 3 mL of saline was injected subcutaneously. During the procedure, the heart was kept moist with drops of 0.9% NaCl. 1-2% lidocaine was delivered epicardially for arrhythmia management as needed. The fiber mesh of the MMBEs implanted in these studies were 7x7 mm and had a predicted mesh density of *ρ*=25% or *ρ*=75% and a CR=0. The adhesive layer was oversized by a few millimeters around the perimeter of the fiber mesh to facilitate MMBE handling and implantation. All implanted MMBEs ranged from 0.06-0.1 mm in thickness (including fiber mesh and adhesive layer).

Upon chest opening, an MI was generated by ligating the LAD about 3-4 mm below the left atria with a suture (7-0 prolene). 30 min after MI, a thin layer of ultrasound gel pad (Aquaflex, Parker labs) was positioned on the epicardial surface. The probe was positioned on top of the gel pad either manually or using a rail system. Saline and ultrasound gel were added if needed to ensure good transmission. A B-mode echocardiogram (300 frame/sec) in the parasternal long axis view (LAX) of the heart was recorded for speckle tracking. Prior to MMBE coupling, the gel pad was removed, and any remaining blood, saline or ultrasound gel in the epicardial surface were cleaned using gauze sponges.

Attaching the MMBEs onto the infarct region after reducing the size of the LV cavity through an inferior vena cava (IVC) occlusion was the most feasible approach to reduce the variations in initial MMBE loading. By temporarily decreasing venous return to the LV, the length of cardiac muscle cells prior to contraction (or preload) is reduced, ‘shrinking’ the LV’s epicardial surface. If MMBEs are attached when the surface area dimensions of the LV are reduced, once the LV returns to its original dimensions, the MMBE will be under a level of tension that is proportional to the percentage of surface area reduction achieved during IVC occlusion. With this approach, if the MMBE dimensions are kept constant and the percentage of volume reduction achieved with an IVC occlusion is similar among animals, the initial level of tension the MMBE will be under will be comparable among animal subjects.

Per this method, the MMBEs were attached onto the epicardial surface using forceps when the LV reached its lowest cavity volume after an IVC occlusion of about 5 s in duration. A corner of the MMBE was pulled to confirm proper adhesion. Following MMBE attachment, a second echocardiogram was acquired. A Vevo 3100 Imaging system with transducer probes MX250S or MX550D was used for all image acquisition. We assumed that our implantation procedure enabled the MMBEs to be coupled under similar levels of initial tension and that similar levels of neurohormonal compensation following MI occurred at the time of MMBE implantation (i.e. 30 min post-coronary ligation). Despite the level of control our platform allows, observing scalable biomechanical changes remains heavily reliant on biological and implantation variability.

The VevoStrain software was used to quantify ventricular epicardial strain in the treatment region from acquired echocardiogram video frames from at least one cardiac cycle. Using the *freecurve* tool, 10 points were placed from base to apex on the epicardial surface of the anterior wall of each animal subject. The points were manually placed from the most basal segment in the frame to the most apical, aiming for the points to be equally spaced and consistent with anatomical landmarks between animals. Longitudinal strain curves of each point during one cardiac cycle were calculated and exported to MATLAB for subsequent analysis.

Two experienced echocardiogram readers identified if the individual points were in a muscle region covered by the attached MMBE (i.e. treatment region) and provided a score of N for no coverage, E for edge of coverage and P for MMBE patch coverage). To capture the nature of contractile muscle motion in the region below the point, readers scored H is the motion was normokinetic, BZ if reduced or hypokinetic, or MI if non-existent or akinetic. Custom MATLAB code was used to calculate the mean longitudinal strain curve, the corresponding standard deviation, and variance of the points marked to be within the patched or treatment region. The longitudinal strain variance of each subject was defined as the area enclosed by the standard deviation of the mean longitudinal strain curve within the treatment region using MATLAB’s *trapz* function. Statistical analysis (paired, nonparametric t-test: Wilcoxon matched-pairs signed rank test) was performed to compare the longitudinal strain variance before and after MMBE placement using GraphPad. A P value < 0.05 was considered to be statistically significant.

## Acknowledgements

The authors would like to thank the Koch Institute Preclinical Imaging Core and Frederick Roberts from FujiFilm VisualSonics, Inc. for their technical assistance, and Dr. Chris Nguyen for providing the deidentified cardiac MRI data file. C.E.V. acknowledges support from the Ford Foundation Predoctoral Fellowship, National Science Foundation Graduate Research Fellowship Program, and the National Institutes of Health (5T32HL007208-45; training grant to the Massachusetts General Hospital Cardiovascular Research Center). D.Q.M. acknowledges support from the Líderes del Mañana Fellowship, the SMA2 Brown Fellowship and the MathWorks Engineering Research Fellowship. J.B. acknowledges support from the SICPA Foundation and Lausanne University Hospital Improvement fund. X.Z acknowledges support from National Institutes of Health (Grants No. 1R01HL153857-01 and No. 1R01HL167947-01), Department of Defense Congressionally Directed Medical Research Programs (Grant No. PR200524P1), and the National Science Foundation (Grant No. 2430106). E.T.R. acknowledges support from the National Science Foundation (CAREER award 1847541) and support from the Institute for Medical Engineering and Science and the Department of Mechanical Engineering at the Massachusetts Institute of Technology.

## Supplementary Figures

**Supplementary Figure 1:**
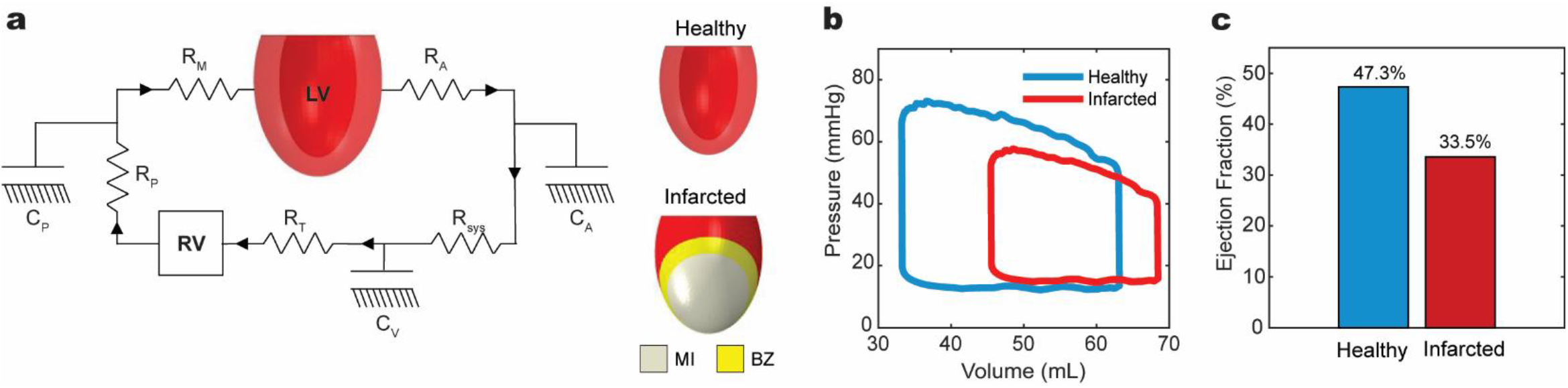
**a)** Lumped parameter circulatory model connected to in silico left ventricle (LV). **b)** Pressure-volume loops generated by healthy and infarcted LV. **c) E**jection fraction calculated from healthy and infarcted LV. (R_M_= mitral valve resistance, R_A_= aortic valve resistance, C_A_= aortic compliance, R_sys_= systemic resistance, C_V_= venous compliance, R_T_= tricuspid resistance, RV= right ventricle, R_P_= pulmonary resistance, C_P_= pulmonary compliance, MI= myocardial infarction, BZ= border zone.

**Supplementary Figure 2:**
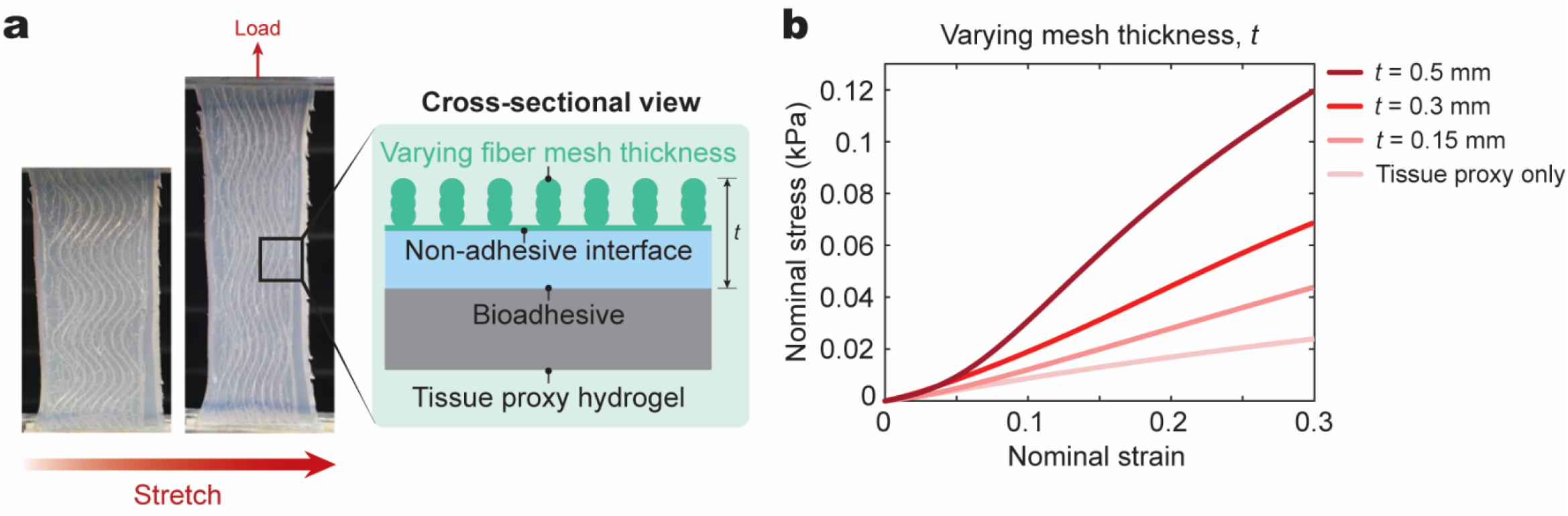
**a)** Image of tissue proxy hydrogel reinforced by a MMBE under uniaxial tensile load. A cross-section of the sample. Increasing the mesh thickness (t) was achieved by printing additional fiber mesh layers. **b)** Uniaxial tensile testing data of tissue proxy hydrogels reinforced by MMBEs of different thickness.

**Supplementary Figure 3:**
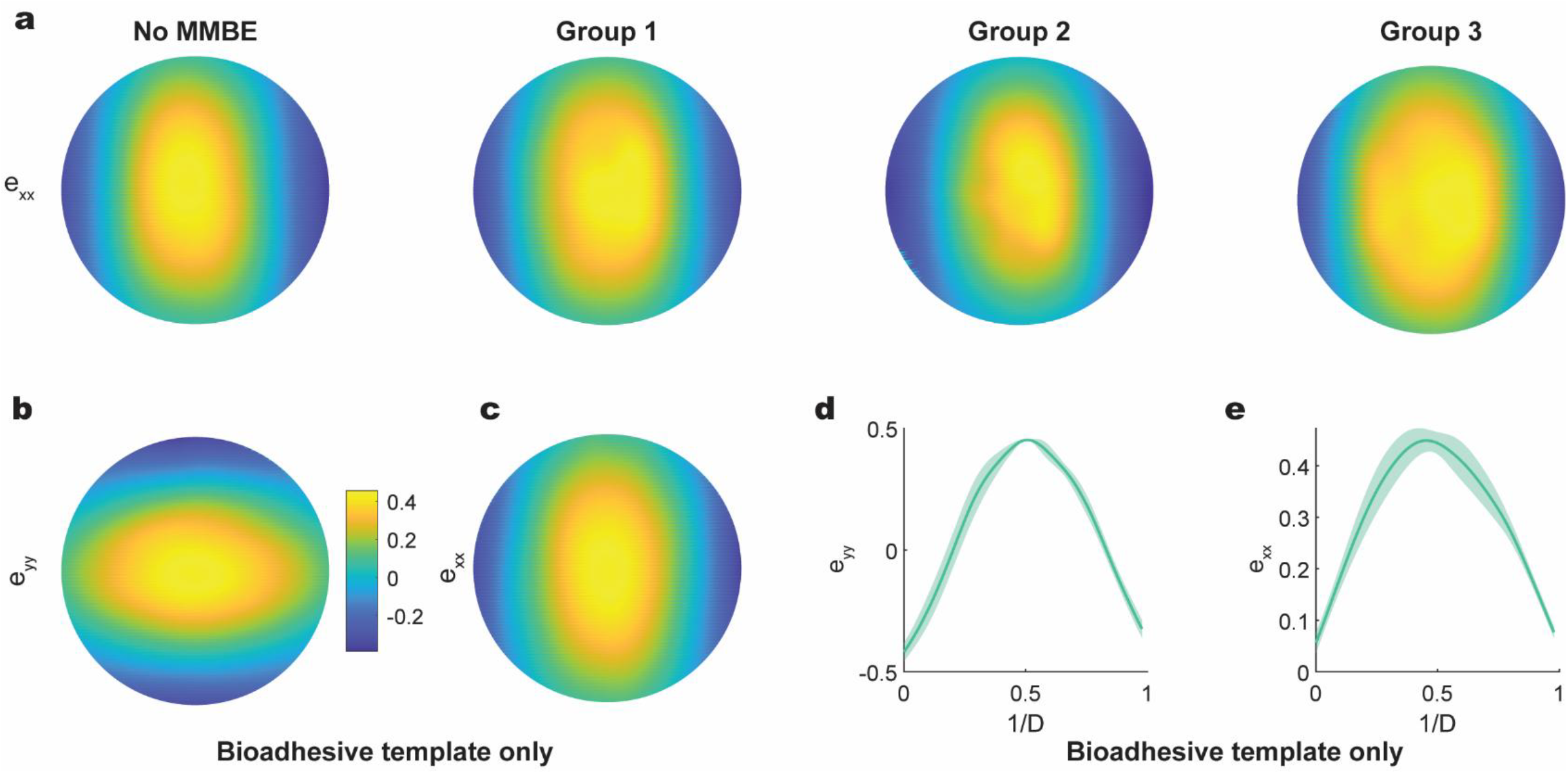
**a)** Representative longitudinal circumferential (exx) strain heat map at 18 mmHg of chamber pressure calculated from digital image correlation (DIC). Representative longitudinal (eyy) **(b)** and circumferential (exx)**(c)** strain heat map at 18 mmHg of chamber pressure calculated from digital image correlation (DIC). (**d**,**e**) Longitudinal and circumferential strain values across the horizontal centerline of the recorded picture frames of width D. n=3 per group; data reported as mean± st.dev.

**Supplementary Figure 4:**
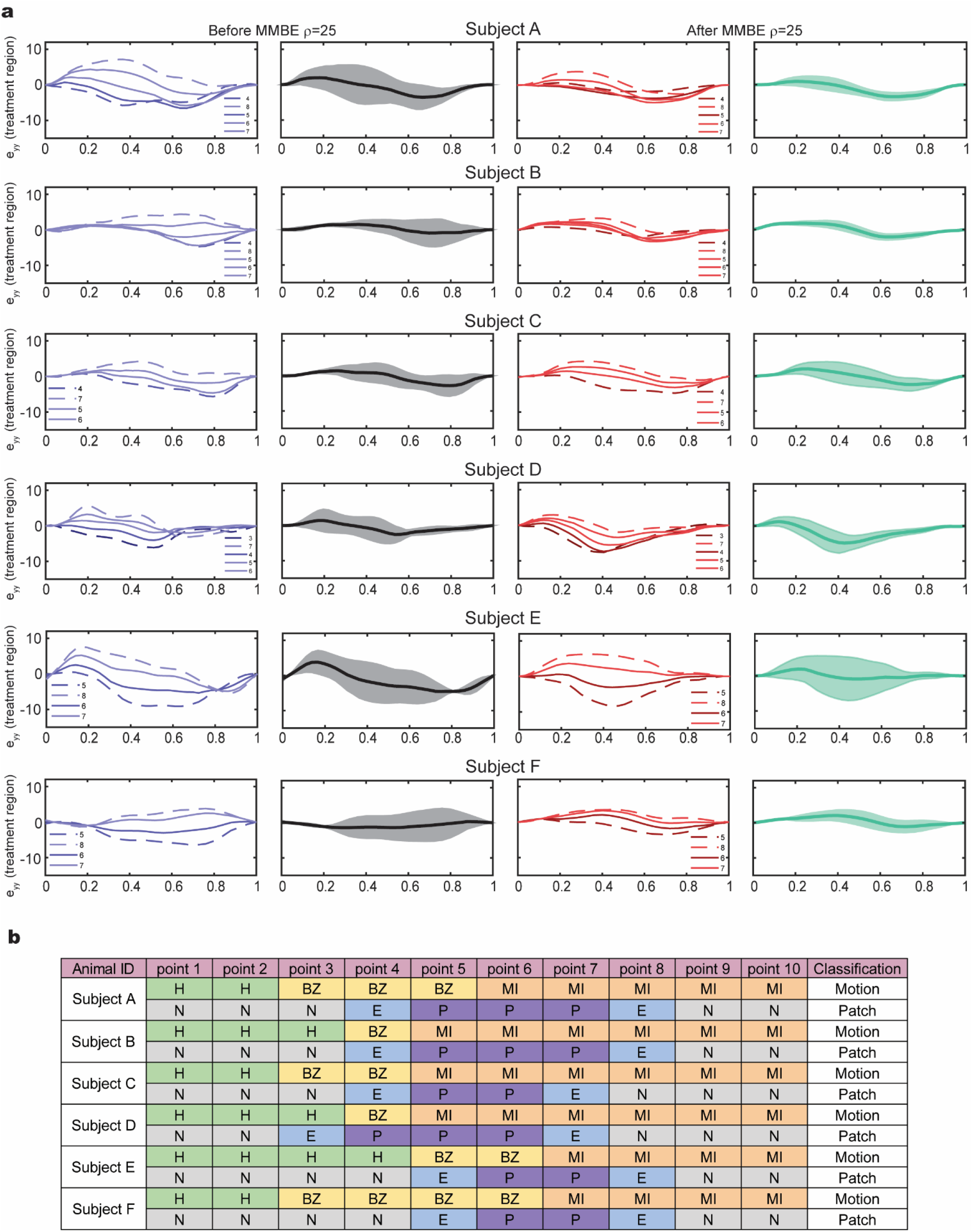
**a)** Individual and mean longitudinal strain curves calculated from each subject before (blue, black) and after (red, green) MMBE reinforcement (design 1; ρ=25) in the treatment region over one cardiac cycle. **b)** Point classification per animal subject in MMBE design 1 group. Contractile muscle motion score: H= normokinesis, BZ= hypokinesis, MI= akinesis. Patch coverage score: N=not covered, E= edge of patch coverage, P= covered by patch.

**Supplementary Figure 5:**
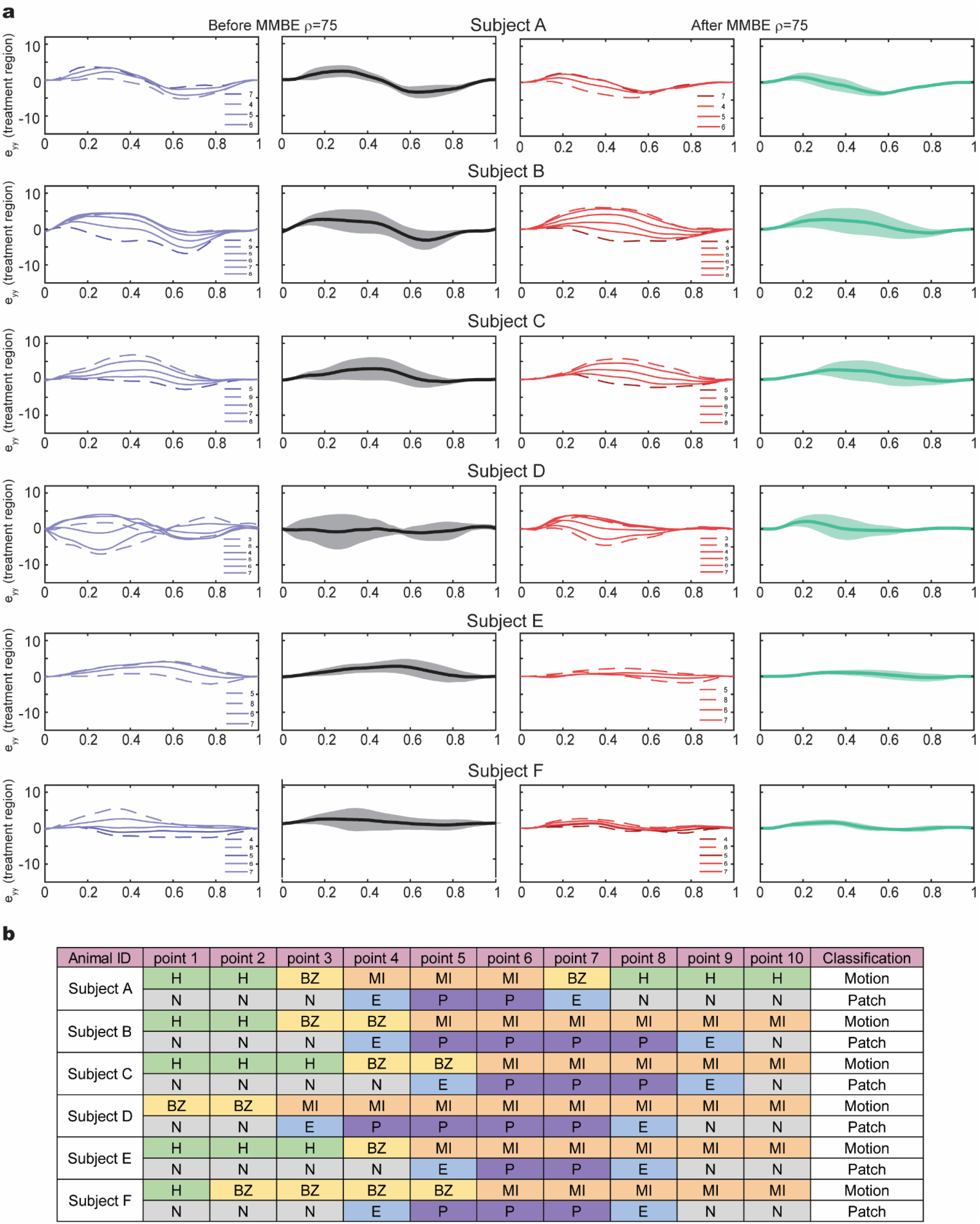
**a)** Individual and mean longitudinal strain curves calculated from each subject before (blue, black) and after (red, green) MMBE reinforcement (design 3; ρ=75) in the treatment region over one cardiac cycle. **b)** Point classification per animal subject in MMBE design 3 group. Muscle motion score: H= normokinesis, BZ= hypokinesis, MI= akinesis. Patch coverage score: N=not covered, E= edge of patch coverage, P= covered by patch.

**Supplementary Figure 7:**
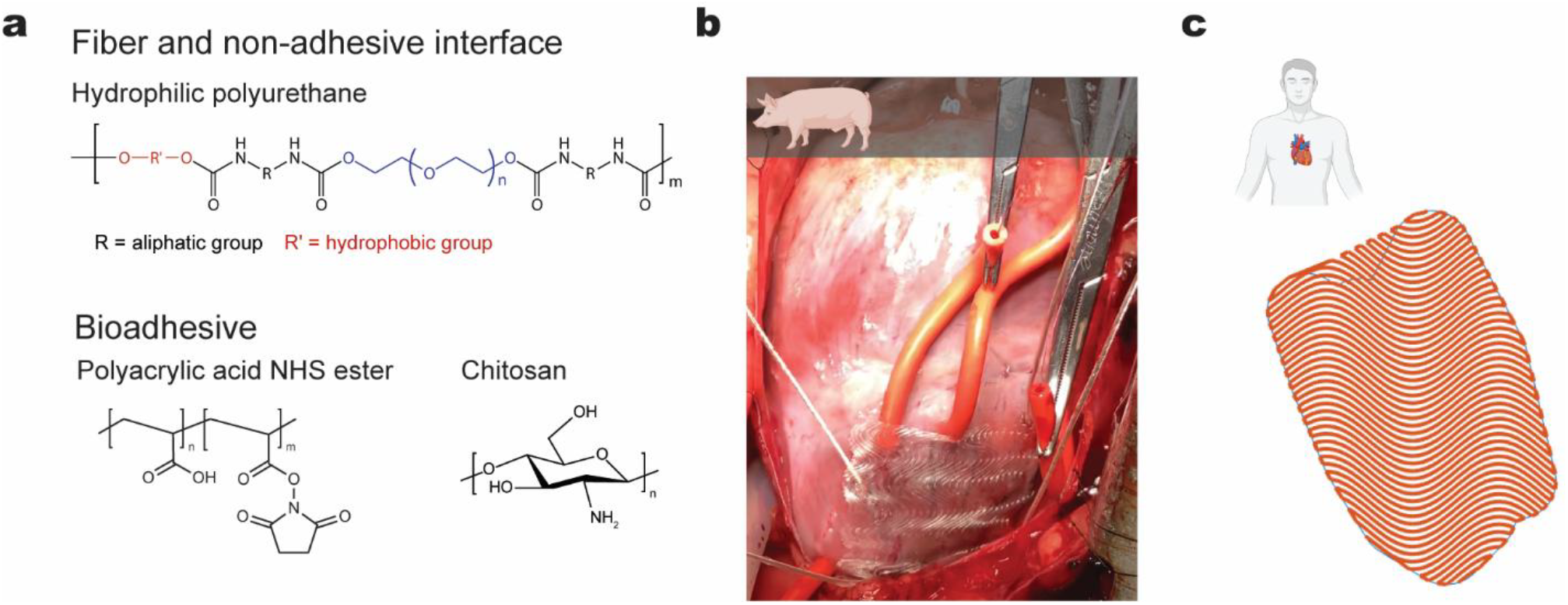
**a)** Chemical compounds of MMBE components. **b)** Sample porcine MMBE fabricated with the developed workflow. **c)** Sample patient-specific MMBE design with circumferentially oriented fibers. The shape of the patient’s infarct region was acquired from a deidentified cardiac MRI dataset.

## References

1. McKay, R. G. et al. Left ventricular remodeling after myocardial infarction: a corollary to infarct expansion. Circulation 74, 693–702 (1986).

2. Cohn, J. N., Ferrari, R. & Sharpe, N. Cardiac remodeling--concepts and clinical implications: a consensus paper from an international forum on cardiac remodeling. Behalf of an International Forum on Cardiac Remodeling. J Am Coll Cardiol 35, 569–582 (2000).

3. Carberry, J., Marquis-Gravel, G., O’Meara, E. & Docherty, K. F. Where Are We With Treatment and Prevention of Heart Failure in Patients Post–Myocardial Infarction? JACC Heart Fail 12, 1157– 1165 (2024).

4. Martin, S. S. et al. 2025 Heart Disease and Stroke Statistics: A Report of US and Global Data from the American Heart Association. Circulation 151, e41–e660 (2025).

5. Jebran, A. F. et al. Engineered heart muscle allografts for heart repair in primates and humans. Nature 2025 639:8054 639, 503–511 (2025).

6. Le, N. T. et al. Stem Cell Therapy for Myocardial Infarction Recovery: Advances, Challenges, and Future Directions. Biomedicines 2025, Vol. 13, Page 1209 13, 1209 (2025).

7. Mathur, A. et al. Five-year follow-up of intracoronary autologous cell therapy in acute myocardial infarction: the REGENERATE-AMI trial. ESC Heart Fail 9, 1152 (2022).

8. Segers, V. F. M. & Lee, R. T. Stem-cell therapy for cardiac disease. Nature 2008 451:7181 451, 937–942 (2008).

9. Huang, K., Hu, S. & Cheng, K. A New Era of Cardiac Cell Therapy: Opportunities and Challenges. Adv Healthc Mater 8, e1801011 (2019).

10. Liu, T. et al. Advanced Cardiac Patches for the Treatment of Myocardial Infarction. Circulation 149, 2002–2020 (2024).

11. Kwon, M. H., Cevasco, M., Schmitto, J. D. & Chen, F. Y. Ventricular restraint therapy for heart failure: A review, summary of state of the art, and future directions. J Thorac Cardiovasc Surg 144, 771-777.e1 (2012).

12. Clarke, S. A., Ghanta, R. K., Ailawadi, G. & Holmes, J. W. Cardiac restraint and support following myocardial infarction. in Studies in Mechanobiology, Tissue Engineering and Biomaterials vol. 15 169–206 (Springer, 2014).

13. Varela, C. E., Fan, Y. & Roche, E. T. Optimizing Epicardial Restraint and Reinforcement Following Myocardial Infarction: Moving Towards Localized, Biomimetic, and Multitherapeutic Options. Biomimetics 2019, Vol. 4, Page 7 4, 7 (2019).

14. Mewhort, H. E. M. et al. Bioactive Extracellular Matrix Scaffold Promotes Adaptive Cardiac Remodeling and Repair. JACC Basic Transl Sci 2, 450–464 (2017).

15. Lin, X. et al. A viscoelastic adhesive epicardial patch for treating myocardial infarction. Nature Biomedical Engineering 2019 3:8 3, 632–643 (2019).

16. Kataoka, A. et al. Application of polymer-mesh device to remodel left ventricular–mitral valve apparatus in ischemic mitral regurgitation. J Thorac Cardiovasc Surg 155, 1485 (2018).

17. Fomovsky, G. M., Clark, S. A., Parker, K. M., Ailawadi, G. & Holmes, J. W. Anisotropic Reinforcement of Acute Anteroapical Infarcts Improves Pump Function. Circ Heart Fail 5, 515 (2012).

18. Estrada, A. C., Yoshida, K., Clarke, S. A. & Holmes, J. W. Longitudinal Reinforcement of Acute Myocardial Infarcts Improves Function by Transmurally Redistributing Stretch and Stress. J Biomech Eng 142, (2020).

19. Clarke, S. A., Goodman, N. C., Ailawadi, G. & Holmes, J. W. Effect of Scar Compaction on the Therapeutic Efficacy of Anisotropic Reinforcement Following Myocardial Infarction in the Dog. J Cardiovasc Transl Res 8, 353–361 (2015).

20. Yuk, H. et al. Dry double-sided tape for adhesion of wet tissues and devices. Nature 2019 575:7781 575, 169–174 (2019).

21. Wu, J. et al. Adhesive anti-fibrotic interfaces on diverse organs. Nature 2024 630:8016 630, 360–367 (2024).

22. Freed, A. D. & Doehring, T. C. Elastic Model for Crimped Collagen Fibrils. J Biomech Eng 127, 587–593 (2005).

23. Gathercole, L. J. & Keller, A. Crimp Morphology in the Fibre-Forming Collagens. Matrix 11, 214– 234 (1991).

24. Ma, Q. et al. A nonlinear mechanics model of bio-inspired hierarchical lattice materials consisting of horseshoe microstructures. J Mech Phys Solids 90, 179–202 (2016).

25. Sack, K. L. et al. Construction and validation of subject-specific biventricular finite-element models of healthy and failing swine hearts from high-resolution DT-MRI. Front Physiol 9, 344767 (2018).

26. Gasser, T. C., Ogden, R. W. & Holzapfel, G. A. Hyperelastic modelling of arterial layers with distributed collagen fibre orientations. J R Soc Interface 3, 15–35 (2006).

27. D’Amore, A. et al. Bi-layered polyurethane - Extracellular matrix cardiac patch improves ischemic ventricular wall remodeling in a rat model. Biomaterials 107, 1–14 (2016).

28. Chen, X., Yuk, H., Wu, J., Nabzdyk, C. S. & Zhao, X. Instant tough bioadhesive with triggerable benign detachment. Proc Natl Acad Sci U S A 117, 15497–15503 (2020).

29. Theocharidis, G. et al. Strain-Programmable Patch for Diabetic Wound Healing. bioRxiv 2021.06.07.447423 (2021) doi:10.1101/2021.06.07.447423.

30. Ashby, M. F. The properties of foams and lattices. Philosophical Transactions of the Royal Society A: Mathematical, Physical and Engineering Sciences 364, 15–30 (2005).

31. Chen, Y., Li, T., Scarpa, F. & Wang, L. Lattice Metamaterials with Mechanically Tunable Poisson’s Ratio for Vibration Control. Phys Rev Appl 7, 024012 (2017).

32. Hadjicharalambous, M., Lee, J., Smith, N. P. & Nordsletten, D. A. A displacement-based finite element formulation for incompressible and nearly-incompressible cardiac mechanics. Comput Methods Appl Mech Eng 274, 213–236 (2014).

33. Holzapfel, G. A. & Ogden, R. W. Constitutive modelling of passive myocardium: A structurally based framework for material characterization. Philosophical Transactions of the Royal Society A: Mathematical, Physical and Engineering Sciences 367, 3445–3475 (2009).

34. Nielles-Vallespin, S. et al. Assessment of Myocardial Microstructural Dynamics by In Vivo Diffusion Tensor Cardiac Magnetic Resonance. J Am Coll Cardiol 69, 661–676 (2017).

35. Guccione, J. M., McCulloch, A. D. & Waldman, L. K. Passive material properties of intact ventricular myocardium determined from a cylindrical model. J Biomech Eng 113, 42–55 (1991).

36. Fan, Y. et al. A comparison of two quasi-static computational models for assessment of intramyocardial injection as a therapeutic strategy for heart failure. Int J Numer Method Biomed Eng 35, (2019).

37. McGarvey, J. R. et al. Temporal Changes in Infarct Material Properties: An In Vivo Assessment Using Magnetic Resonance Imaging and Finite Element Simulations. Annals of Thoracic Surgery 100, 582–589 (2015).

38. Ambrosi, D., Arioli, G., Nobile, F. & Quarteroni, A. Electromechanical Coupling in Cardiac Dynamics: The Active Strain Approach. https://doi.org/10.1137/100788379 71, p605–621 (2011).

39. Guccione, J. M., Waldman, L. K. & McCulloch, A. D. Mechanics of actiwe contraction in cardiac muscle: Part II—cylindrical models of the systolic left ventricle. J Biomech Eng 115, 82–90 (1993).

40. Pilla, J. J., Gorman Iii, J. H. & Gorman, R. C. Theoretic Impact of Infarct Compliance on Left Ventricular Function. (2009) doi:10.1016/j.athoracsur.2008.11.044.

41. Guan, D., Yao, J., Luo, X. & Gao, H. Effect of myofibre architecture on ventricular pump function by using a neonatal porcine heart model: From DT-MRI to rule-based methods. R Soc Open Sci 7, (2020).

42. Wenk, J. F. et al. First evidence of depressed contractility in the border zone of a human myocardial infarction. Annals of Thoracic Surgery 93, 1188–1193 (2012).

